# Humans sacrifice decision-making for action execution when a demanding control of movement is required

**DOI:** 10.1101/2020.04.08.028936

**Authors:** Amélie J. Reynaud, Clara Saleri Lunazzi, David Thura

## Abstract

A growing body of evidence suggests that decision-making and action execution are governed by partly overlapping operating principles. Especially, previous work proposed that a shared decision urgency/movement vigor signal, possibly computed in the basal ganglia, coordinates both deliberation and movement durations in a way that maximizes the reward rate. Recent data support one aspect of this hypothesis, indicating that the urgency level at which a decision is made influences the vigor of the movement produced to express this choice. Here we investigated whether conversely, the motor context in which a movement is executed determines decision speed and accuracy. Twenty human subjects performed a probabilistic decision task in which perceptual choices were expressed by reaching movements toward targets whose size and distance from a starting position varied in distinct blocks of trials. We found strong evidence for an influence of the motor context on most of the subjects’ decision policy but contrary to the predictions of the “shared regulation” hypothesis, we observed that slow movements executed in the most demanding motor blocks in terms of accuracy were often preceded by faster and less accurate decisions compared to blocks of trials in which big targets allowed expression of choices with fast and inaccurate movements. These results suggest that decision-making and motor control are not regulated by one unique “invigoration” signal determining both decision urgency and action vigor, but more likely by independent, yet interacting, decision urgency and movement vigor signals.

**NEW & NOTEWORTHY:** Recent hypotheses propose that choices and movements share optimization principles derived from economy, possibly implemented by one unique context-dependent regulation signal determining both processes speed. In the present behavioral study conducted on human subjects, we demonstrate that action properties indeed influence perceptual decision-making, but that decision duration and action vigor are actually independently set depending on the difficulty of the movement executed to report a choice.

## INTRODUCTION

Animals, including humans, are faced with decisions about actions on a daily basis, and they behave to seek rewards while avoiding punishments and minimizing energy expenditure. Because the evaluation of reward, risk, and effort governs our action choices, investigating how the brain processes these variables is critical to improve our understanding of adapted or dysfunctional goal-directed behavior.

Importantly, the subjective value of a given activity is not only limited to its related reward, risks, and efforts. It also depends on the amount of time invested in it, as time strongly discounts the value of rewards (Myerson and Green, 1995). Therefore, what is ultimately most adaptive is to choose options that maximize one’s global reward rate (Bogacz et al., 2010; Balci et al., 2011), which occurs when the decision and action processes are sufficiently accurate but not overly effortful and time-consuming. As a consequence, nearly all decision scenarios present decision-makers with speed-accuracy-effort trade-offs during both decision-making and action execution, and the brain must control both processes to maximize the rate of reward.

Because trade-offs during decision and action have been typically studied in isolation, mechanisms allowing a coordinated maximization of reward rate are still elusive. Recent promising advances suggest, however, that motor control and choices, including economic ones, are governed by partly overlapping optimization principles (Shadmehr et al., 2010, 2019; Haith et al., 2012; Choi et al., 2014; Yoon et al., 2018; Carland et al., 2019). First, human and non-human primates move faster and with a shorter reaction time toward items that they value more (Kawagoe et al., 1998; Summerside et al., 2018; Revol et al., 2019). Second, humans take motor costs into account during both motor (Cos et al., 2011, 2012, 2014; Morel et al., 2017) and non-motor (Burk et al., 2014; Marcos et al., 2015; Diamond et al., 2017; Hagura et al., 2017) decisions and effortful reaches impose a cost for decision-making similar to cost functions in motor control (Wickler et al., 2000; Shadmehr et al., 2016; Morel et al., 2017; Reppert et al., 2018). Finally, in the foraging paradigm where one makes decisions regarding how long to stay and accumulate reward from one patch, and then moves with certain speed to another patch, the goods collection duration and the vigor (movement speed and duration) with which human subjects move from one reward site to another are governed by a mechanism allowing to maximize the overall capture rate (Yoon et al., 2018).

In line with this shared optimization hypothesis, we and others have proposed that control of urgency is critical for reward rate maximization during decision-making between actions (Ditterich, 2006; Churchland et al., 2008; Standage et al., 2011; Thura et al., 2012; Malhotra et al., 2017, 2018). Urgency is a context-dependent, motor-related signal that grows over the time course of deliberation, pushing the decision-related neural activity toward the commitment threshold (Thura and Cisek, 2014; Kira et al., 2015; Murphy et al., 2016; Steinemann et al., 2018). Remarkably, we demonstrated in a changing evidence decision task that urgency level at decision time strongly influences speed and duration of the following motor commands: early decisions, usually made based on strong sensory evidence but low urgency, were followed by long movements (in terms of duration) whereas late decisions, relying on weak sensory evidence but strong urgency, were followed by faster movements. Then, when subjects were encouraged to make faster and less accurate decisions in distinct blocks of trials, movements were faster compared to blocks encouraging slow and accurate choices. These results imply that a shared invigoration signal, possibly computed in the basal ganglia, coordinates the unified adaptation of the speed-accuracy trade-off during both decision-making and action execution in order to control the rate of reward (Thura et al., 2014; Thura and Cisek, 2016, 2017; Cisek and Thura, 2018; Thura, 2020).

We proposed a model of this hypothetical mechanism, labeled the “shared regulation” hypothesis (Figure 1A, Thura et al., 2014). In this model, speed-accuracy trade-offs for deciding and acting are influenced by a shared decision urgency/movement vigor signal. As a consequence, the context-dependent urgency level at which a decision is made should determine the vigor (duration and speed scaled by amplitude) of movements produced to express this choice and conversely, the context-dependent vigor of movements executed to express a choice should predict the level of urgency with which that choice is made. Recent behavioral and neurophysiological data collected in both trained monkeys and naïve humans strongly support the former prediction (Thura et al., 2014; Thura and Cisek, 2016; Thura, 2020). The latter prediction, namely whether or not the fastest choices are made in motor contexts encouraging the most vigorous movements (Figure 1B), remains, however, to be tested.

**Figure 1:**
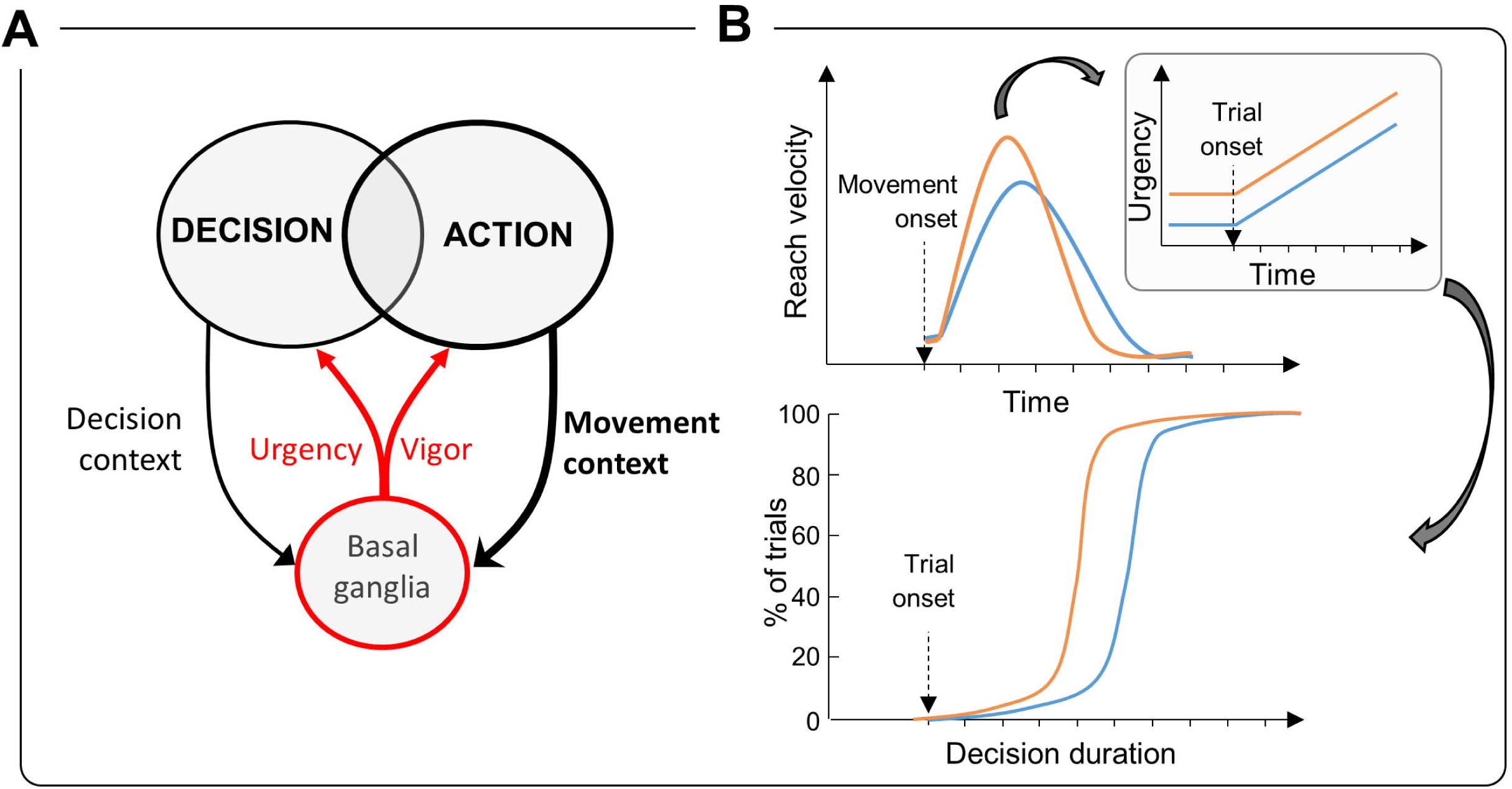
The “shared regulation” hypothesis. A. Simplified hypothetical mechanism of a shared regulation of decision and movement durations by one unique invigoration (decision urgency/movement vigor) signal, possibly computed in the basal ganglia (Thura et al., 2014; Thura and Cisek, 2017). The thick black lines illustrate the manipulation of the motor context, tested in the present study, leading to the modulation of the urgency/vigor signal. B. The shared regulation hypothesis makes a simple prediction regarding the effect of the motor context in which a decision is made on the duration of that decision: if a context encourages execution of vigorous (faster, shorter) movements (orange) to report choices, then the urgency level in this context should be raised compared to another context in which movements need to be less vigorous but more accurate (blue). As a consequence, equally difficult decisions made in the vigorous block of trials should be on average shorter than those made in the block encouraging slow and accurate movements.

To this aim, we conducted an experiment in which human subjects performed a probabilistic decision task in which perceptual choices were expressed by reaching movements toward targets whose size and distance from the starting point varied across blocks of trials, allowing us to assess the effects of the motor context on subjects’ decision policy. In the present work, the speed and duration of the movements are considered as indicators of action vigor and the movement speed-accuracy trade-off are used to modulate this vigor.

## MATERIALS AND METHODS

### Participants

Twenty-three healthy, human subjects (ages: 18-41; 17 females; 21 right-handed) participated in this study. All gave their consent orally before starting the experiment. The ethics committee of Inserm (IRB00003888) approved the protocol on March 19^th^, 2019. Each participant was asked to perform two experimental sessions. They received monetary compensation (20 € per completed session) for participating in this study. Among them, twenty (ages: 20-41; 16 females; 18 right-handed) completed at least two sessions and have thus been included in the present dataset.

### Dataset

The decision and motor behaviors of most of the subjects (17/20) have been described in a recent publication aimed to report the effect of decision strategy on movement properties in human subjects (Thura, 2020). This analysis showed that according to the shared regulation hypothesis, the urgency level at the time of decision commitment strongly influences movement kinematics, with urgency-based decisions leading to vigorous movements. In the present paper, we analyzed data of the same subjects along with data from 3 additional ones, but we grouped trials depending on movement constraints (target size/movement amplitude configurations, see below), allowing us to test on the same subjects the reverse side of the shared regulation hypothesis, i.e. the effects of motor context on decision policy.

### Setup

The subjects sat in an armchair made planar reaching movements using a handle held in their dominant hand (Figure 2A). A digitizing tablet (GTCO CalComp) continuously recorded the handle horizontal and vertical positions (100 Hz with 0.013cm accuracy). Target stimuli and cursor feedback were projected by a DELL P2219H LCD monitor (60 Hz refresh rate) onto a half-silvered mirror suspended 26 cm above and parallel to the digitizer plane, creating the illusion that targets floated on the plane of the tablet. Unconstrained eye movements and pupil area of a subset of subjects were recorded using an infrared camera (ISCAN, sampling rate of 120 Hz, data not shown).

**Figure 2:**
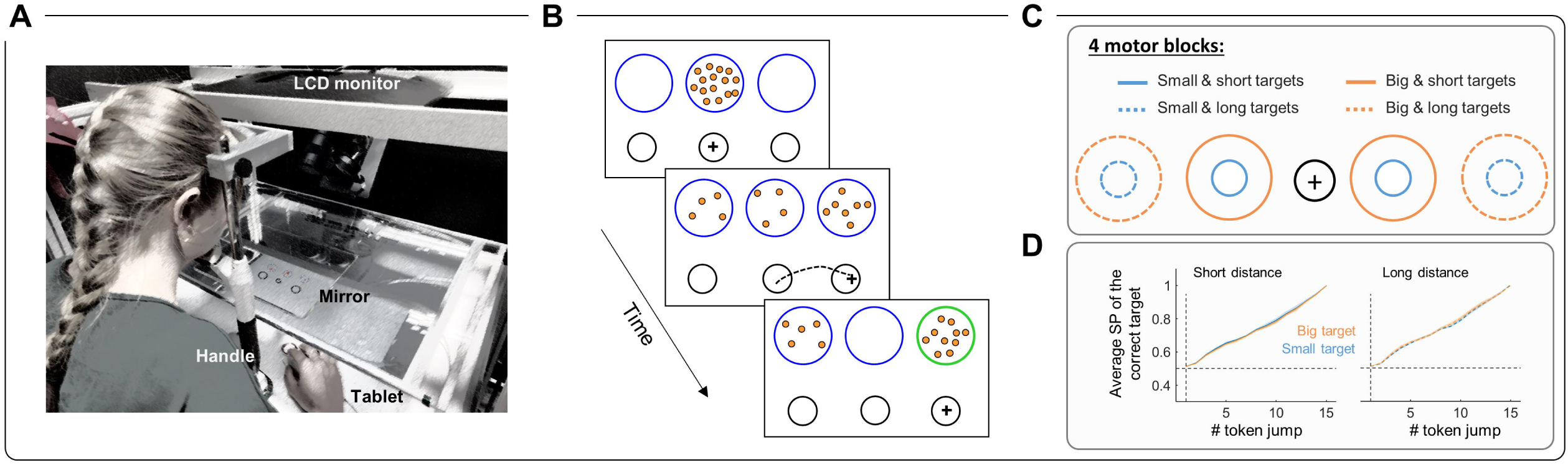
Apparatus, experimental design, and conditions. A. Experimental apparatus. B. Time course of a trial in the tokens task. C. Motor conditions, i.e. movement target size and distance combinations. In distinct blocks of trials, both lateral targets could be either small and located close to the starting circle (black), small and located far from the starting circle, big and located close to the starting circle or big and located far from the starting circle. D. Average success probability profiles of trials experienced by subjects in each of the four motor conditions.

### Tasks

The subjects performed a modified version of the tokens task (Figure 2B, see Cisek et al., 2009 for the original version). They were faced with a visual display consisting of three blue circles (1.5 cm radius) placed horizontally at a distance of 6 cm of each other (the “decision” stimuli). In the central blue circle, 15 small tokens were randomly arranged. Positioned 12 cm below, three black circles, organized horizontally as well defined the “movement” stimuli. While the central black circle radius was kept constant at 0.75 cm, the size of the two lateral black circles and their distance from the central circle could vary, set to either 0.75 (small) or 1.5 cm (big) of radius, and either 6 (short) or 12 cm (long) of distance from the central circle, in distinct blocks of trials. This design allowed us to define four motor blocks depending on the size/distance combination of the two targets: “small/short”, “small/long”, “big/short” and “big/long” (Figure 2C).

A trial was initiated when the subject moved and hold the handle into the small black central circle (starting position) for 500ms. Tokens then started to jump, one by one, every 200ms in one of the two possible lateral blue circles. The subjects’ task was to decide which of the two lateral blue circles would receive the majority of the tokens at the end of the trial. They reported their decisions by moving the handle into the lateral black circle corresponding to the side of the chosen blue circle. Importantly, subjects were allowed to make and report their choice at any time between the first and the last jump. Arm movement duration could not exceed 800ms, irrespective of the motor block. If a movement exceeds 800ms (too slow) or if it reaches the target but fails to stop in it within 800ms (inaccurate), the trial is considered as a movement error trial. Once the choice is properly reported, the remaining tokens jumped more quickly to their final circles. In separate blocks of trials, this post-decision interval was set to either 20ms (“fast” decision block) or 150ms (“slow” decision block). The acceleration of the remaining tokens implicitly encouraged subjects to decide before all tokens had jumped into their respective lateral circles, to save time and increase their rate of reward. Note that each reaching movement carries a temporal cost with respect to reward rate maximization (see equation 3) because the remaining tokens accelerate only when action is completed. After holding the handle in the target for 500ms, visual feedback about decision success or failure (the chosen decision circle turning either green or red, respectively) was provided after the last token jump. A 1500ms period (the inter-trial interval) preceded the following trial.

Before and after the tokens task described above, each subject also performed 100 trials of a delayed reach task (DR task). This task was identical to the tokens task except that there was only one lateral decision circle displayed at the beginning of the trial (either at the right or the left side of the central circle with 50% probability) and all tokens moved from the central circle to this unique circle at a GO signal occurring after a variable delay (1000 ± 150ms). They executed 2 different motor blocks of 25 trials each before the tokens task and the 2 other motor blocks (25 trials each) after the tokens task. This DR task was used to estimate the sum of the delays attributable to sensory processing of the stimulus display as well as to response initiation in each motor condition.

### Instructions

In a given session, subjects were asked to complete one slow decision block and one fast decision block of the tokens task. To complete a decision block (either fast or slow), subjects had to make 160 correct choices, indirectly motivating them to optimize successes per unit of time. After the first block was completed, a short break was offered to the subject. Within each decision block, the size of the movement targets and their distance from the starting circle, i.e. the motor blocks, were varied every 40 trials. In a session, each motor block was thus performed twice, once in the slow decision block, and once in the fast decision block.

Subjects performed two sessions (test-retest design), one a day, and each of them separated by a maximum of 7 days. In session #1 subjects always started the tokens task in the slow decision block with the following succession of motor blocks: small/short, small/long, big/short, and big/long; followed by the execution of the fast decision block with the same motor blocks order. To prevent any block-related confounding effect, the order of decision and motor blocks presentation was reversed in session #2. Before the first session, we explicitly described to the subjects the principle of each decision block, specifying that deciding quickly in the fast block was more advantageous in terms of time-saving than in the slow block (because of the larger acceleration of the remaining tokens) but that such hasty behavior could also lead to more erroneous decisions. A short recall was provided before starting the second session. Because subjects were informed that they had to complete a given number of correct responses in each session, they were all aware that they were presented with a speed/accuracy trade-off in this task. A practice period consisting of performing 20 tokens task trials in the slow decision and big/short motor blocks was proposed at the beginning of the first session, mainly allowing subjects to get familiar and comfortable with the manipulation of the handle on the tablet. Among the 23 subjects who participated in this study, two have been tested six and seven times. The additional sessions performed by these two subjects are not described in the present report.

### Data analysis

All arm movement data were analyzed off-line using MATLAB (MathWorks). Reaching characteristics were assessed using the subjects’ movement kinematics. Horizontal and vertical position data were first filtered using a tenth-degree polynomial filter and then differentiated to obtain a velocity profile. Onset and offset of movements were determined using a 3.75 cm/s velocity threshold. Peak velocity was determined as the maximum value between these two events and endpoint error was defined as the Euclidian distance separating the target center from the movement endpoint location. The dispersion of movement end-points is visualized with confidence ellipses representing an iso-contour of the Gaussian distribution, defining the region that contains 95% of all samples in each condition.

We computed at each moment during a trial the success probability *p*_*i*_(*t*) associated with choosing each target *i*. For a total of 15 tokens, if at a particular moment in time the right target contains *N*_*R*_ tokens, whereas the left contains *N*_*L*_ tokens, and there are *N*_*C*_ tokens remaining in the center, then the probability that the target on the right will ultimately be the correct one (i.e., the success probability of guessing right) is as follows:

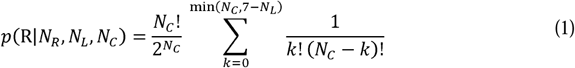

To characterize the success probability profile of each trial, we calculated this quantity (with respect to either the correct target or the target ultimately chosen by the subject, depending on purposes) for each token jump. To ensure that the difficulty of decisions was homogeneous among subjects and experimental conditions, we controlled the sequence of trials experienced by subjects in each session. Especially, we interspersed among fully random trials (20% of the trials in which each token is 50% likely to jump into the right or the left lateral circle) three special types of trials characterized by particular temporal profiles of success probability. Subjects were not told about the existence of these trials. 30 % of trials were so-called “easy” trials, in which tokens tended to move consistently toward one of the circles, quickly driving the success probability *p*_*i*_(*t*) for each toward either 0 or 1. Another 30% of trials were “ambiguous”, in which the initial token movements were balanced, making the *p*_*i*_(*t*) function close to 0.5 until later in the trial. The last special trial type was called “misleading” trials (20%) in which the 2-3 first tokens jumped into the incorrect circle and the remaining ones into the correct circle. In all cases, even when the temporal profile of success probability of a trial was predesigned, the actual correct target was randomly selected on each trial. Importantly, the sequence of trials was designed such as the proportion of each trial type was similar in each decision and motor condition (Figure 2D).

To estimate the time at which subjects committed to their choice (decision time, DT) on each trial in the tokens task, we detected the time of movement onset, defining the subject’s reaction time (RT) and subtracted from it her/his mean sensory-motor delays (SM) estimated based on her/his reaction times in the same motor block of the delayed reach task performed the same day. Decision duration (DD) was computed as the duration between the DT and the first token jump. Equation 1 was then used to compute for each trial the success probability at the time of the decision (SP).

Calculation of subjects’ accuracy criterion at decision time relies on the available sensory evidence at that time. Because it is very unlikely that subjects can calculate Equation 1, we computed a simple “first-order” approximation of sensory evidence as the sum of log-likelihood ratios (SumLogLR) of individual token movements as follows (Cisek et al., 2009, page 11567, provides more details on this analysis):

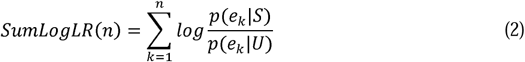

where *p*(*e*_*k*_|*S*) is the likelihood of a token event *e*_*k*_ (a token jumping into either the selected or unselected target) during trials in which the selected target *S* is correct, and *p*(*e*_*k*_| *U*) is its likelihood during trials in which the unselected target *U* is correct. The SumLogLR metric is thus proportional to the difference in the number of tokens that have moved in each circle before the moment of decision. To characterize the decision policy of a given subject in a given block of trials, we binned trials as a function of the total number of tokens that moved before the decision and calculated the average SumLogLR for each bin.

To quantify subjects’ performance relative to the task objective, i.e. complete a given number of correct decisions, assuming they tried to complete each block as quickly as possible, we first calculated for correct and bad decisions the reward rate (RR), using a local definition (Haith et al., 2012; Thura et al., 2012) which corresponds to the expected number of correct choices per unit of time. This is computed as follows:

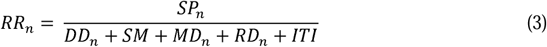

where SP_*n*_ is the probability that the choice made on trial *n* was correct, DD_*n*_ is the time taken to make the decision, SM is the sensorimotor delays (specific to each motor context but constant for a given session), MD_*n*_ is the movement duration, RD_*n*_ is the duration of the remaining token jumps after the target is reached, and ITI is the inter-trial interval (fixed at 1500ms). Then from the average reward rate computed in each motor block, we calculated the average number of correct choices per minute and deducted from it the time necessary to complete a given number of correct choices in each condition of interest.

Comparisons of decision duration, success probability, movement duration, peak velocity, accuracy, and block duration between conditions performed for each subject are statistically tested with Wilcoxon-Mann-Whitney (WMW, two-sided rank-sum) tests. The effect of motor condition on sensory evidence at decision time as a function of decision duration is statistically tested with analyses of covariance (ANCOVAs). For these analyses, very fast decisions made before token jump #4 are discarded. Decisions made before jump #4 were rare (see Thura, 2020) and success probability homogeneity (if subjects decide before token jump #4 it is likely because the first three tokens jumped into the same target) at that time makes data exclusion reasonable. The proportion of inadequate movements in small target conditions (small/short and small/long blocks) is statistically compared to the proportion of inadequate movements in big target conditions (big/short and big/long blocks) for each subject with chi-square tests. For all statistical tests, the significance level is set a 0.05.

## RESULTS

### Effect of motor context on motor behavior in the tokens task

As expected, the motor context in which decisions were reported strongly influenced subjects’ movement properties and performance. First, we calculated the percentage of trials in which an inadequate movement was performed to express a choice, i.e. a movement exceeding 800ms (too slow) or failing to stop and maintain the position in the target within 800ms (inaccurate). In the first session, most subjects (18/20) performed significantly more inadequate movements in the small target (small/short and small/long blocks) condition compared to the big target (big/short and big/long blocks) condition (Chi-square tests, p < 0.05). Movement “error” rates within blocks are the following across the population: small/long target blocks:18.8% ± 6.8; small/short: 5.5% ±3.1; big/long: 4.5% ± 2.7; big/short: 1% ±1.3. Despite an overall slight decrease, the same impact of motor constraint was observed on movement error rate during session #2: 19 out of 20 subjects made more inadequate movements in the small target compared to the big target condition (Chi-square tests, p < 0.05), with the following error rates in each of the four motor contexts: small/long target blocks: 16.7% ±4.5; small/short: 4.8% ±1.8; big/long: 2.35% ±1.5; big/short: 1.1% ±1.3). Figure 3A shows the dispersion of movement endpoints in one example subject who performed the tokens task in the four motor blocks. In this plot, correct and inadequate (too slow or inaccurate) movements trials are included. Confidence ellipses (containing 95% of all samples in each condition) largely extend outside of movement targets in small target trials, especially when targets are far from the starting center, whereas they almost entirely fit into movement targets in big target trials.

**Figure 3:**
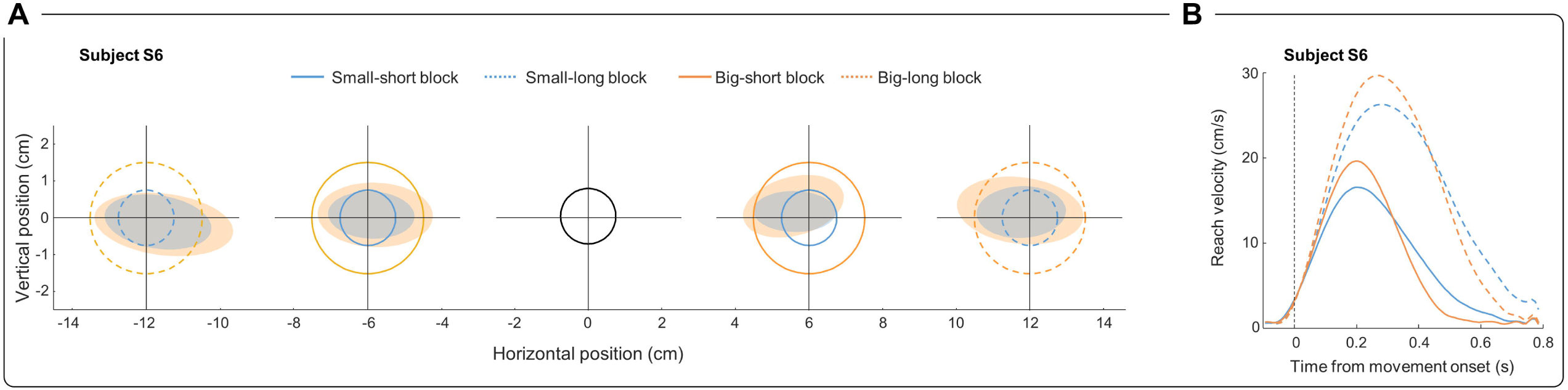
Motor behavior in one example subject. A. Panel shows the motor visual display depicted in Figure 2, along with shaded ellipses illustrating for each motor condition and side with respect to the start circle (black) the dispersion (an iso-contour of the Gaussian distribution) of one example subject reaching endpoints in the tokens task. Each ellipse contains 95% of the data in each condition, and trials include correct and inadequate (too slow or inaccurate) movements executed in the two sessions and the two decision conditions (slow and fast). B. Reach velocity profiles of the same subject in the four motor conditions. Same color/style convention as in A. Only adequate movements are included.

Then, we focused analyses on trials in which an adequate movement was performed to express a choice, irrespective of the outcome of that choice. As expected, reaching movement properties, in terms of velocity peak, duration, and endpoint “error” (the distance between target center and movement offset location) were affected by the motor context in which movements were executed. Figure 3B shows for the same representative subject the reaching velocity profiles in trials sorted as a function of the four motor blocks. Unsurprisingly, movement velocity was largely higher and duration longer in long target (dotted lines) compared to short target trials (solid lines), regardless of the size of the target. The size of the target also modulated movement speed and duration but to a lesser extent. Movements were indeed slightly faster and shorter when executed toward big targets (orange lines) compared to those executed toward small targets (blue lines).

This effect of motor context on movement properties was observed on the vast majority of subjects performing either the tokens or the delayed reach (DR) task. To simplify comparisons in the following analyses, we grouped trials depending on (1) target size, defining two conditions, small versus big target conditions, regardless of target distance from the starting circle, and (2) target distance from the starting circle, defining two other conditions, short versus long target conditions, regardless of target size.

First, most of the subjects reported decisions by making significantly faster (15 out of 20 subjects, WMW test, p<0.05), shorter, in terms of duration (17 out of 20 subjects, WMW test, p<0.05) and more dispersed (18/20, WMW test, p<0.05) movements in the big target compared to the small target condition (figure 4A). Second, all subjects reached long targets with significantly faster and longer movements compared to movements executed toward short targets (WMW test, p<0.05, figure 4B, left and middle panels). In this distance contrast, we observed that endpoint distances from target center were not as consistently modulated as in the size contrast, being significantly larger for the long target compared to the short target condition in only 9 out of 20 subjects (WMW test, p<0.05, figure 4B, right panel). The same influence of target characteristics on reaching velocity, duration, and accuracy was found in the DR task (not shown). Finally, the influence of target characteristics on movement parameters was similar in the two experimental sessions and the two decision blocks (slow and fast, not shown).

**Figure 4:**
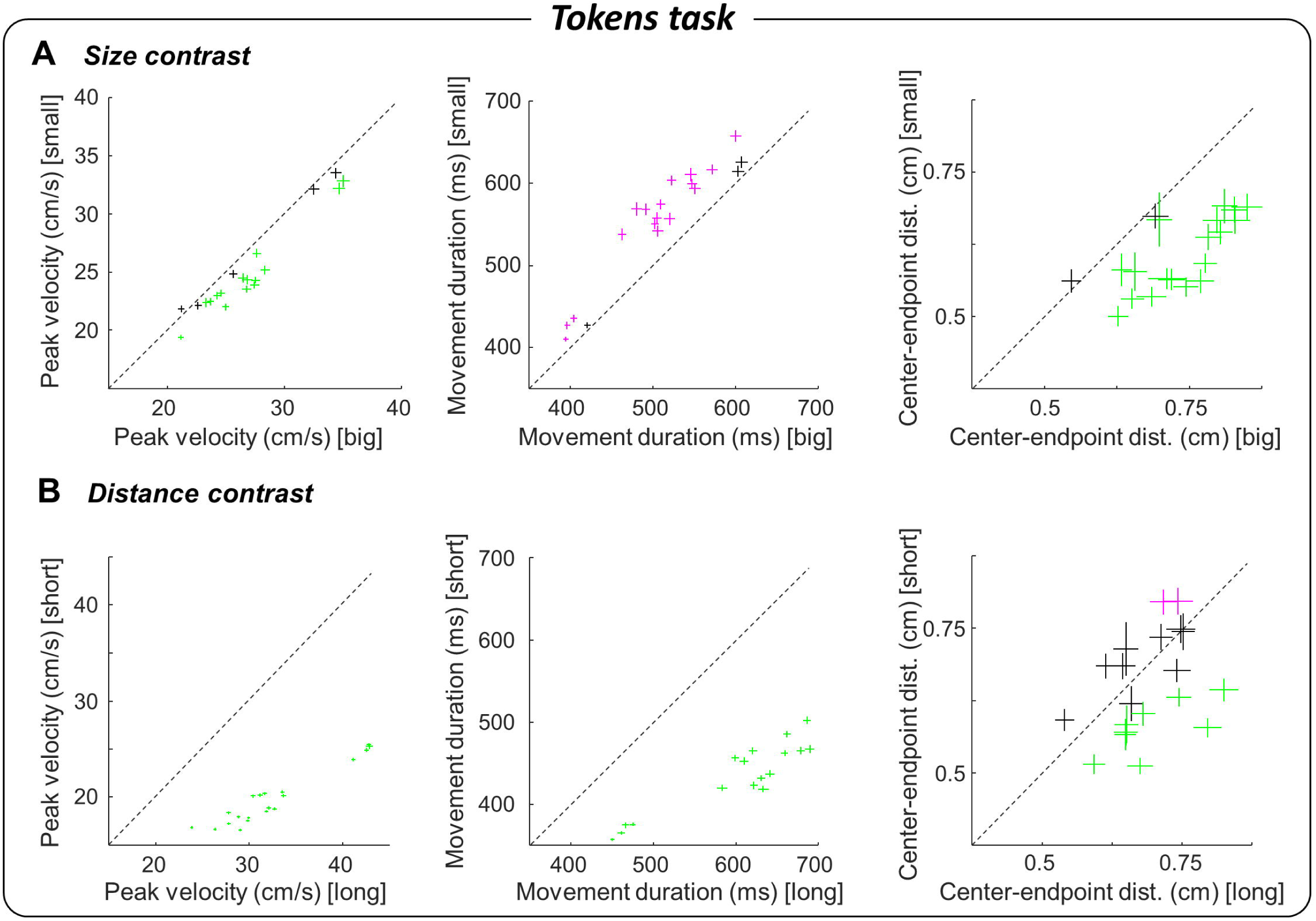
Effect of motor context on population motor behavior. A. Average reaching movement peak velocity (left), duration (middle) and target center-endpoint distance (right) of each subject during big target (big/short and big/long blocks, x-axis) and small target (small/short and small/long blocks, y-axis) conditions performed in the tokens task. Green (magenta) pluses indicate the mean and SE for subjects for whom data is larger (smaller) in the big target condition compared to the small target condition and the difference was significant (WMW test, p < 0.05). Data include trials collected from both sessions #1 and #2, in both the slow and fast decision blocks. B. Same as A for trials executed in the long target (small/long and big/long blocks, x-axis) versus the short target (small/short and big/short blocks, y-axis) condition.

To summarize, manipulating the target characteristics in distinct blocks of trials successfully modulated reaching movement properties, encouraging subjects to either emphasize speed or accuracy to execute movements in these blocks to express their choices. In the following section, we assess whether or not these context-dependent adjustments of motor parameters influenced the decision policy leading to the actions executed to report choices.

### Effect of motor context on subjects’ decision behavior

To determine the potential impact of movement context on decision policy, we first analyzed subjects’ decision duration (regardless of the decision outcome) by sorting trials depending on target characteristics, irrespective of the session and the decision condition (slow or fast). By first comparing decisions made in big (big/short and big/long) versus small (small/short and small/long) target trials, we found that decisions were overall shorter in the small target compared to the big target condition (1099 versus 1154ms). Importantly, the difference is significant for half of the population (WMW test, p<0.05, figure 5A, left panel). Only one subject behaved the opposite way, making significantly faster choices when allowed to report them with fast, less accurate reaching movements. Importantly, we found virtually no difference between the average decision difficulties (quantified as success probability profiles, see Methods and figure 2D) in the two motor conditions, excluding a role of the sensory evidence experienced by the subjects in the difference of decision duration observed between small and big target contexts. Did this shortening of decision duration affect choice accuracy? To answer that question, we analyzed the amount of sensory evidence that subjects needed to commit to their choices (i.e. their accuracy criterion, computed as the sum of the log-likelihood ratios, see Methods), as a function of decision duration for the two motor conditions, small and big target trials (Figure 5A, middle panel). First, the level of sensory evidence that subjects required before committing to a choice decreased as a function of decision duration, irrespective of motor conditions (ANCOVA, SumLogLR, time effect, F_(1,347)_ = 164, p < 0.0001). This observation suggests that the more time is elapsing over the time course of a trial, the more decisions rely on a sensory-agnostic signal. In our previous studies as well as in others, this decreasing accuracy criterion is interpreted as a behavioral signature of an urgency-gating mechanism of decision-making, which in short describes the decision variable as the combination of sensory evidence with an urgency signal and the decision is made when the decision variable reaches a constant threshold (Cisek et al., 2009; Thura et al., 2012).

**Figure 5:**
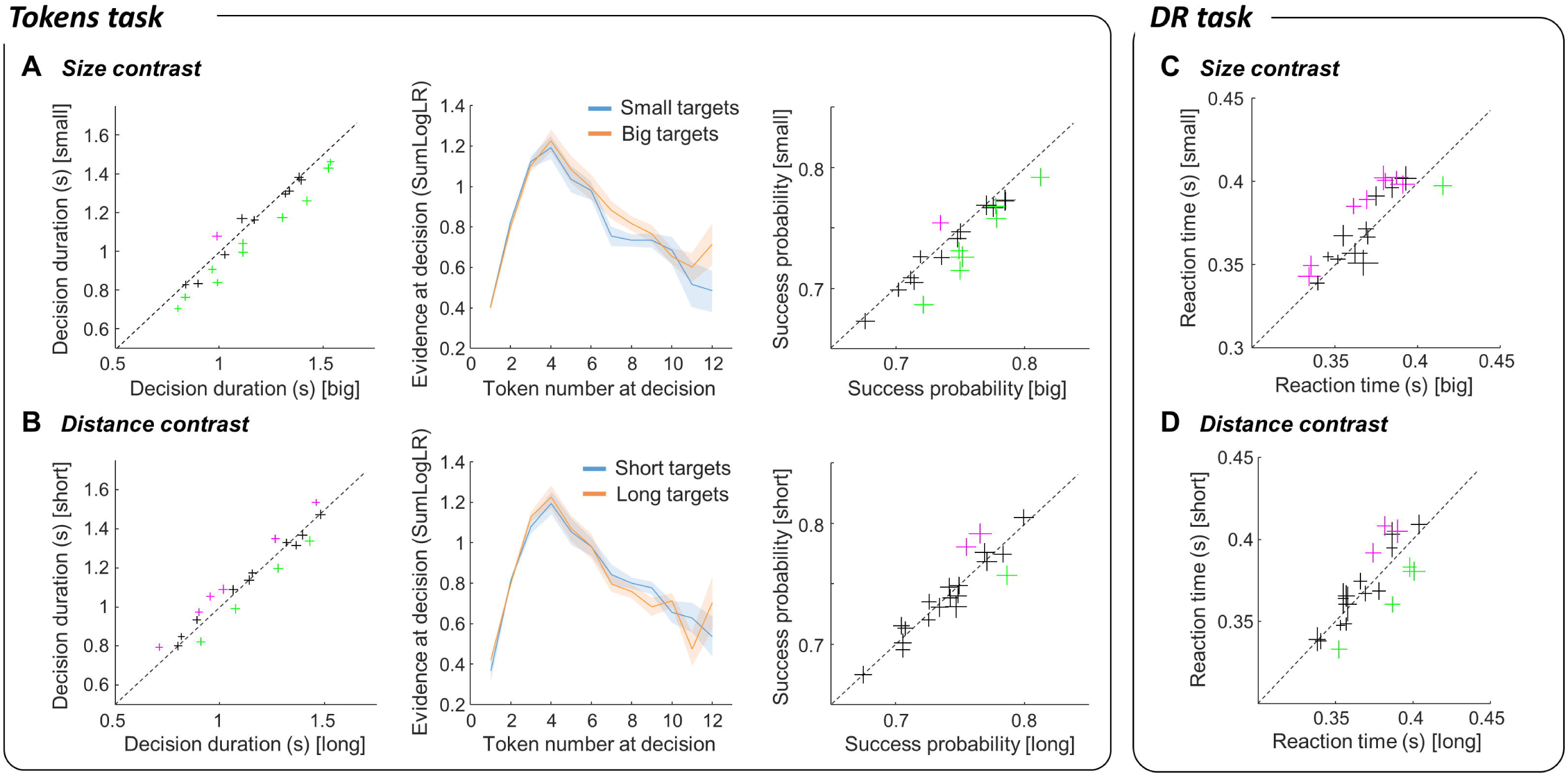
Effect of motor context on decision behavior. A. Left: Average decision duration of each subject during big (x-axis) and small (y-axis) target conditions performed in the tokens task. Same convention as in Figure 4. Middle: Average (± SE) evidence at decision time across subjects as a function of decision duration in the small (blue) and the big (orange) target conditions of the tokens task. Right: Mean success probability of each subject during big (x-axis) and small (y-axis) target conditions performed in the tokens task. Same convention as in Figure 4. Data include trials collected from both sessions #1 and #2, in both the slow and fast decision blocks. B. Same as A for trials executed in the long versus short target conditions. C. Average reaction time of each subject during big (x-axis) and small (y-axis) target conditions performed in the delayed reach task. D. Same as C for trials executed in the long (x-axis) versus the short (y-axis) target condition.

Importantly for the present report, we found that the accuracy criterion of subjects performing the tokens task in small target trials was significantly lower than in big target trials, for any decision made after token jump #3 (ANCOVA, SumLogLR, target size effect, F_(1,347)_ = 4.63, p = 0.03). This indicates that subjects were more willing to tolerate less sensory evidence to make their choices in small target compared to big target trials. As a consequence, decisions were usually less likely to be correct in the small target compared to the big target context (Figure 5A, right panel). This decrease of success probability in small target trials was significant in 7 out of 10 subjects showing a significant decrease of decision duration as a function of target size (WMW test, p<0.05).

We next compared decision durations in short versus long target trials, a contrast that strongly modulates movement speed of all subjects (Figure 4B, left panel). We found that the impact of target distance, and thus movement speed, on decision duration was less consistent at the population level compared to the impact of target size described above (Figure 5B, left panel). Indeed, we observed that 6 subjects made significantly longer decisions in the short target compared to the long target condition (WMW test, p<0.05), 4 subjects behaved the opposite way (WMW test, p<0.05), and the 10 remaining ones did no behave differently, in terms of decision duration, between the two motor conditions. We also found that target distance did not significantly influence the quantity of sensory information used by subjects to commit to their choice (ANCOVA, SumLogLR, target size effect, F_(1,346)_ = 0.13, p = 0.72, Figure 5B, middle panel), and the success probability of these choices was only rarely significantly modulated as a function of target distance (Figure 5B, right panel).

We next analyzed the effect of target size and distance on subjects’ reaction times (RT) in the delayed reach (DR) task. In the DR task, no volitional commitment needed to be made as subjects were instructed with both the correct target and when to execute their response (see Methods). In this task, we found that subjects’ RTs were overall longer in small target compared to big target trials (375 versus 367ms), with a significant difference for 8 out of 20 subjects (WMW test, p<0.05), and only one subject behaving significantly the opposite way (Figure 5C). Interestingly, we found a significant correlation between the modulation of decision duration by target size in the tokens task and the modulation of reaction time in the same conditions in the DR task. In other words, the more subjects expedited decisions in the small target condition of the tokens tasks, the more they slowed down their response initiation in the same condition in the DR task (Pearson correlation, r = -0.495, p = 0.026, Figure 6). By contrast, reaction times were less homogeneously affected by the distance condition in the DR task. Four subjects reacted faster in short compared to long target blocks, and 3 subjects behaved the opposed way (WMW test, p<0.05, Figure 5D).

**Figure 6:**
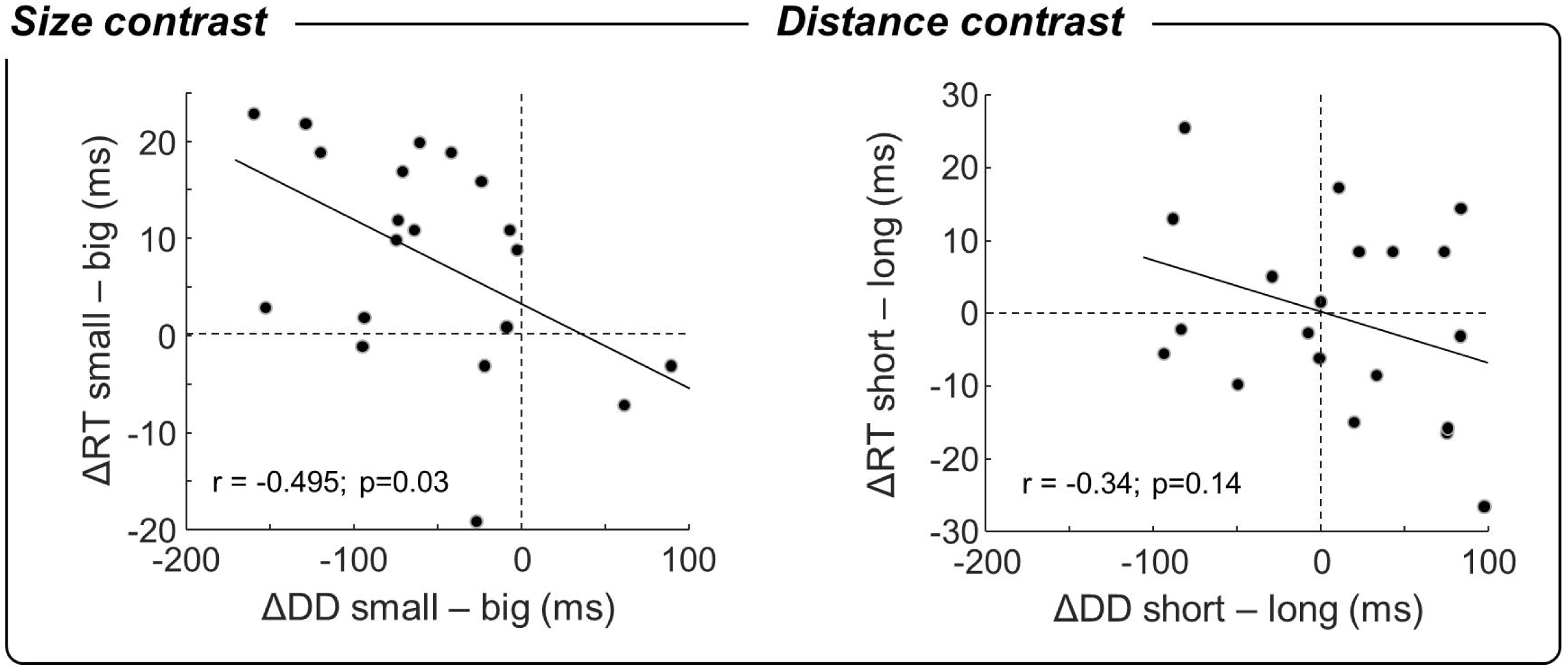
Relationship between the effect of motor context on the decision and instructed tasks. Left: Correlation between the difference of decision duration in small versus big target conditions in the tokens task (x-axis) and the difference of reaction time in the same conditions in the delayed reach task (y-axis). Each dot shows data from one individual subject. Right: Same as Left for the distance contrast (short versus long target conditions).

To assess whether the effect of target size on decision policy was dependent on the decision context, i.e. the slow or fast decision blocks of the tokens task, we computed subjects’ decision duration, success probability, and sensory evidence at decision time for each of the two size conditions, separately for the two decision blocks. In a recent report (Thura, 2020), we describe in detail subjects’ behavior in the two decision conditions. Quickly, the “slow” decision block of trials encourages slow and accurate decisions because the tokens that remain in the central decision circle after movement completion accelerate only a little compared to the pre-decision period (see Methods).

By contrast, in the “fast” block of trials, the remaining tokens accelerate a lot, allowing subjects to potentially save a lot of time by deciding quickly, permitting to eventually maximize their reward rate. In Thura, 2020 we showed that subjects behaved accordingly, making faster (1028 vs 1229ms across subjects) and less accurate (0.87 versus 0.97) decisions in the fast block compared to the slow block of trials (see the average distributions of decision duration across subjects in the

two decision blocks in Figure 7A,B). In the present report, we demonstrate that the impact of target size on decision policy, especially accuracy, is larger in the slow block than in the fast block of trials. Indeed, decision durations were significantly modulated by target size in 8 out of 20 subjects performing the slow block whereas they were modulated in only 6 subjects performing the tokens task in the fast condition (WMW test, p<0.05). Moreover, the accuracy criterion was significantly higher for big target compared to small target trials in the slow block (ANCOVA, SumLogLR, size effect, F_(1,345)_ = 13.6, p = 0.0003) but not in the fast block (F_(1,298)_ = 0.1, p = 0.75, Figure 7A,B, left panels). As a consequence, success probability was strongly influenced by target size in the slow block (significantly modulated in 9 out of 20 subjects, WMW test, p<0.05) whereas effects were more balanced in the fast blocks (Figure 7A,B, right panels).

**Figure 7:**
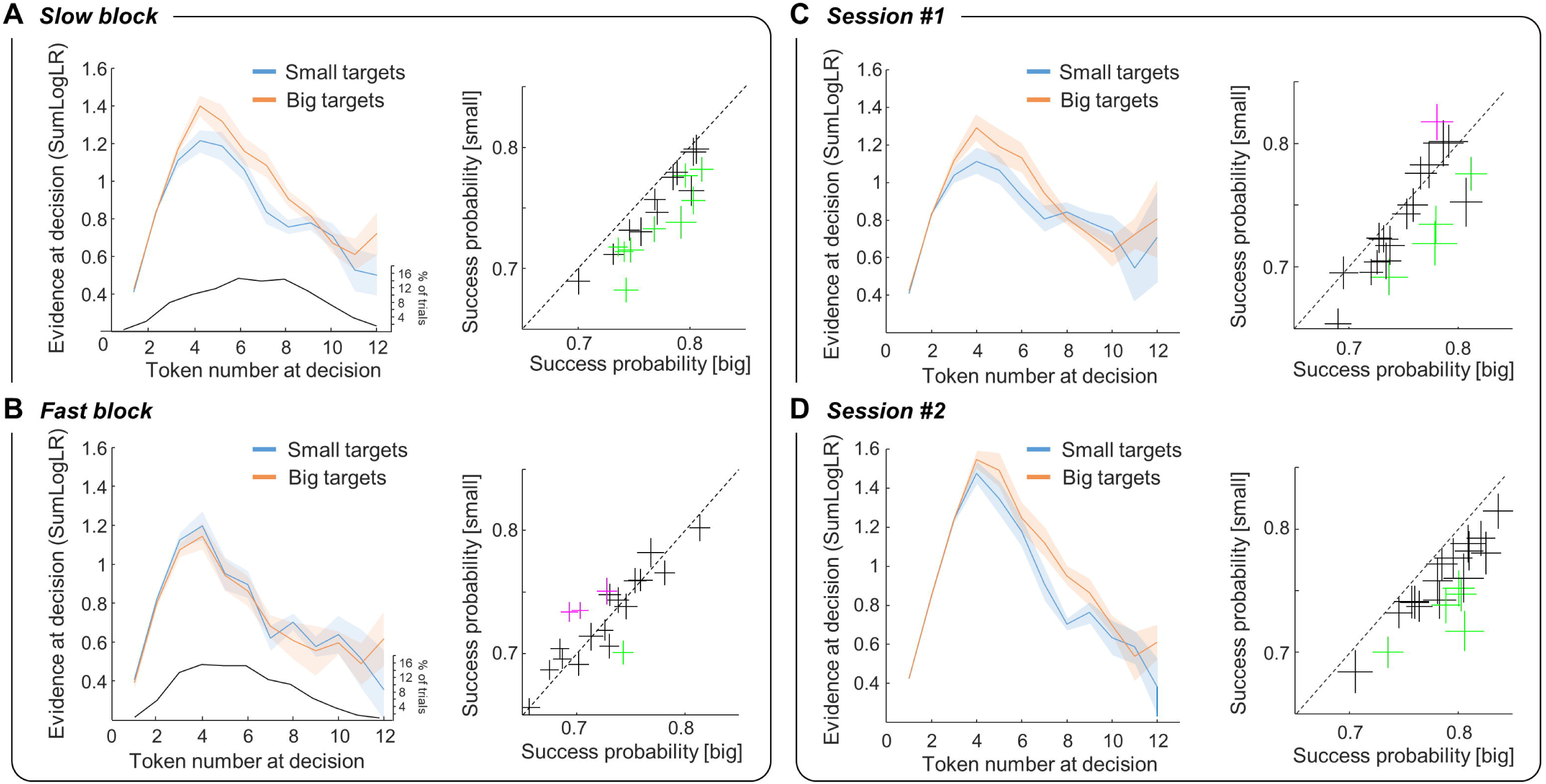
Effect of motor context on decision accuracy depending on decision context and experience. A. Left: Average (± SE) evidence at decision time across subjects as a function of decision duration in small (blue) and big (orange) target conditions performed in the “slow” decision block of the tokens task. The black line below shows the average distribution of decision duration across subjects in the slow block. Right: Average success probability of each subject during big (x-axis) and small (y-axis) target conditions performed in the slow decision block of the tokens task. Data from both sessions #1 and #2 are included. Same convention as in Figure 4. B. Same as A for decisions made in the “fast” decision block of the tokens task. C. Same as A for decisions made in the first session, including only slow decision blocks. D. Same as C for decisions made during the second session.

Next, we analyzed the effect of target size on decision policy depending on the level of experience of subjects in the tokens task. To do this, we computed subjects’ decision duration, success probability, and sensory evidence at decision time for decisions made in the slow decision block for each of the two target size conditions, separately for the two experimental sessions. Overall, we found that the impact of the target size on decision policy did not strongly evolve with training. Decision durations were slightly more modulated by target size in the first session than in the second sessions (5/20 and 3/20 subjects with a significant effect of target size on decision duration in session #1 and #2, respectively; WMW test, p<0.05), but accuracy criterion (ANCOVA, SumLogLR, size effect, F_(1,320)_= 2.5, p=0.1 in session #1; F_(1,330)_ = 10.5, p=0.0013 in session #2) and to a lesser extent, success probability (4/20 and 5/20 subjects with a significant effect of target size on decision duration in session #1 and #2, respectively; WMW test, p<0.05) were more affected by target size in session #2 compared to session #1 (Figure 7C,D).

Finally, we evaluated the impact of the faster and less accurate choices in the small target condition compared to the big target condition on subjects’ performance in the tokens task. Because it has been shown that subjects seek to optimize their rate of correct responses rather than their absolute accuracy (Balci et al., 2011), performance is estimated as the duration that subjects needed to complete each motor block. Thus, by calculating the rate of reward and deducting from it the amount of time necessary to complete the different motor blocks in each session (see Methods), we found that this duration was significantly longer in the small target condition compared to the big target condition across subjects, regardless of the session performed, when subjects performed the tokens task in the slow decision block (WMW test, p = 0.0013, Figure 8, left panel). By contrast, we found no significant difference in block duration between small and big target conditions in the fast decision block of trials (WMW test, p = 0.11, Figure 8, right panel).

**Figure 8:**
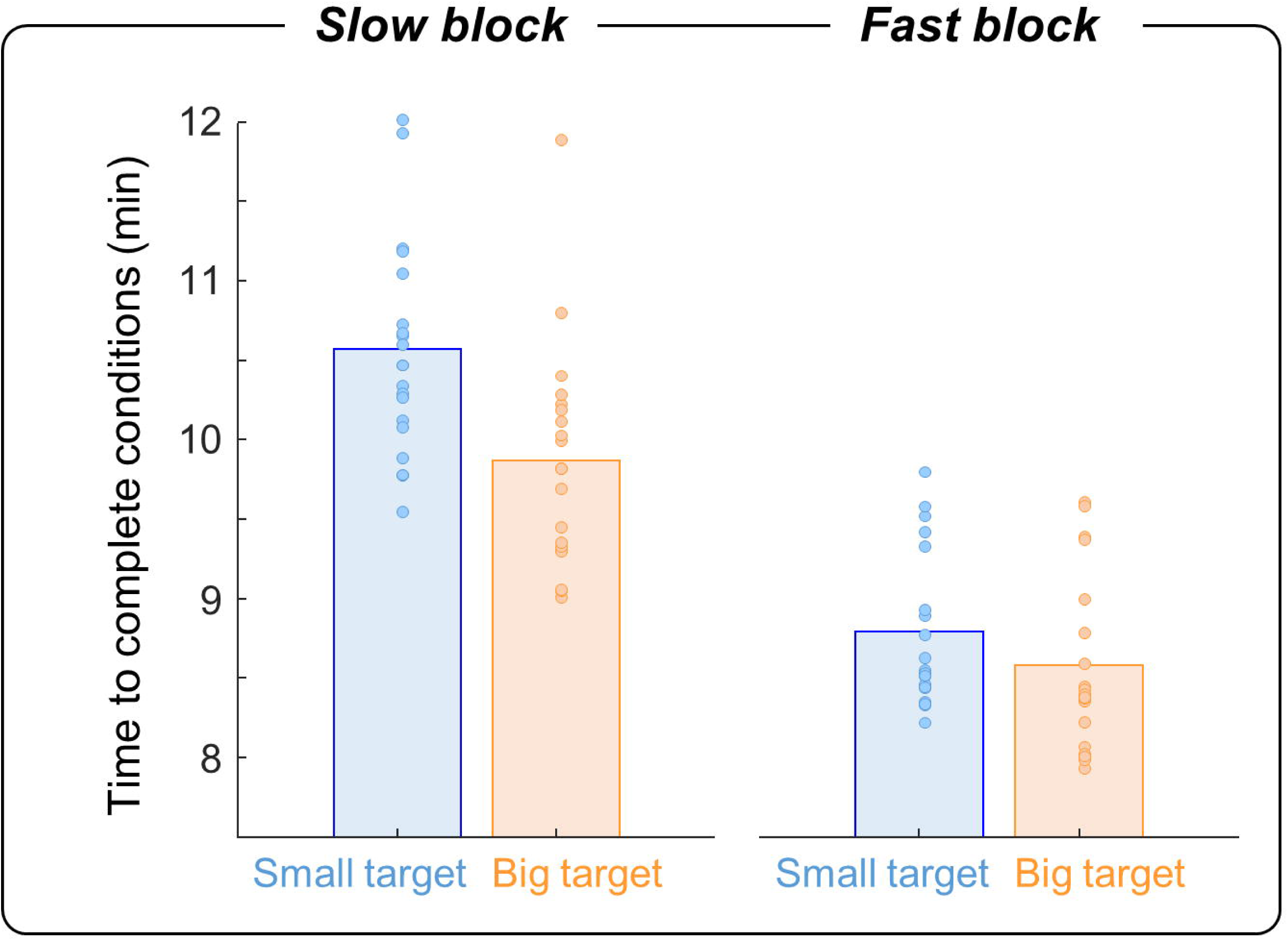
Influence of target size on the expected duration of blocks. Bars show the average expected time necessary to complete a block of 80 trials, computed based on reward rate in each condition, in the small (blue) and big (orange) target block across subjects and sessions, in the slow (left) and fast (right) decision block of trials. Dots illustrate individual data.

## DISCUSSION

In this study, we assessed whether the motor context in which perceptual decisions between actions are made influences human subjects’ decision strategy, as predicted by the recently proposed “shared regulation” hypothesis (Thura et al., 2014). This model conceives decision and action as a continuum, regulated by unspecific signals. As a consequence, a motor context favoring vigorous movements should be preceded by fast decisions because of the activation of one unique invigoration signal possibly computed in the basal ganglia (Cisek and Thura, 2018). We found that motor context indeed often influences decision-making but contrary to the prediction of the shared regulation hypothesis, decisions preceding slow and accurate actions were faster, rather than slower, compared to decisions made in blocks allowing more vigorous and less accurate actions.

In the present task, the action vigor (indicated by movement speed and duration) is assumed to be determined depending on the speed-accuracy trade-off of each motor condition. However, motivation factors, such as the movement energetic cost, may have contributed to shape action vigor as well (Mazzoni et al., 2007). With the present design, we cannot disentangle the contribution of the accuracy and energy costs on vigor definition, the slower movements executed toward the small targets being also less energetically costly. However, if we assume that the less energetically costly movements should increase the subjects’ implicit motivation to decide and act (Mazzoni et al., 2007), those movements should be executed faster than the effortful ones. Yet, our data indicate the opposite results. We thus believe that the accuracy requirement is the main factor that determined movement vigor in our experiment.

### Motor costs influence motor and perceptual decision-making

The present results first add to the many recent observations that challenge the classic view of behavior organization, inherited from cognitive psychology, in which perception, decision, and action are considered as temporally separate and serial processes (Pylyshyn, 1984). Indeed, in ecological scenarios, sensory or value-based decisions are very often expressed by actions that are themselves associated with risks and costs. For instance, a monkey deciding between reaching toward a grape or a nut may prefer the nut but time and energy expenditure associated with opening its shell may rather encourage him to go for the grape. Because it has been extensively demonstrated that the brain tends to control behavior in such a way that the expected value of a choice is maximized while all types of cost are minimized (Neumann and Morgenstern, 1944; Todorov and Jordan, 2002; Gold and Shadlen, 2007; Christopoulos and Schrater, 2015; Christopoulos et al., 2015; Diamond et al., 2017), any potentially penalizing factor, including motor costs, should influence the perceptual judgment leading to a potential reward.

In the past decade, several studies have demonstrated that motor costs influence decision-making when choices only rely on movement properties (i.e. motor decisions). Cos and colleagues showed that when humans make rapid choices between reaching actions, they tend to choose the one that carries the lowest biomechanical cost (Cos et al., 2011, 2014). Morel and colleagues found that biomechanics affects action selection too, but among duration, amplitude, direction and force, they observed that movement duration is perceived as the greatest cost by subjects (Morel et al., 2017). Finally, Michalski and colleagues observed that movement amplitude and direction influence the probability of switching from one ongoing movement to another in a common real-life scenario where one has to decide while already acting (Michalski et al., 2020).

Other work addressed the effects of motor costs on decision-making beyond purely motor choices, i.e. when the decision primarily relies on perceptual or value information, as in the present work. In three of these experiments using the random dots motion discrimination task, data indicate an effect of motor constraints on non-motor decision-making. Burk and colleagues demonstrated that physical effort affects the proportion of changes of mind made by subjects during the deliberation period: the more the change of mind requires a significant energetic cost, the less subjects are willing to perform it (Burk et al., 2014). Another study showed that asymmetric biomechanical cost biases perceptual decisions, with subjects more systematically choosing targets associated with movements of lower cost, even if these choices were detrimental to accuracy (Marcos et al., 2015). In agreement with this observation, Hagura and colleagues demonstrated that motion discrimination is influenced by the physical resistance applied to the response. Intriguingly, they showed that motor costs also bias vocally-expressed judgments, suggesting that actions changed how subjects perceived the stimuli themselves (Hagura et al., 2017). It is important to note that in these three studies, each of the two potential targets was assigned a specific motor cost during a given choice. By contrast, in the present work, the two targets were always associated with the same motor cost, and that cost was varied between blocks of trials. The present report is thus to our knowledge the first to show that the motor context in which a movement is performed influences the strategy of subjects during decision-making.

### A flexible mechanism for regulating decision and movement durations

Decisions about actions typically include a period of deliberation that ends with the commitment to a choice, which then leads to the overt expression of that choice through action execution, at the end of which the reward can be at last consumed. Because decision and action processes are so inextricably linked, it is natural to imagine that they could at least partly share operating principles to maximize the utility of behavior. Decision and action could indeed be considered as a continuum during which regulation signals would affect both processes agnostically, in a unified manner. In agreement with this hypothesis, it has been proposed that movement selection, preparation, and execution are parameterized following economical rules, varying depending on utility estimation: high valued options lead to faster reaction times and movement speed, and high-perceived effort discount option’s value, leading to slower reaction and longer movements (Kawagoe et al., 1998; Wickler et al., 2000; Shadmehr et al., 2010, 2016, 2019; Haith et al., 2012; Choi et al., 2014; Morel et al., 2017; Reppert et al., 2018; Summerside et al., 2018; Yoon et al., 2018; Revol et al., 2019).

Our previous results support this hypothesis of a coordination between decision and action durations during behavior. For instance, within fixed decision and motor contexts, both humans and monkeys shorten their movement duration in trials in which decision duration is prolonged, as if extended deliberation duration was compensated by increasing the action speed so that the next opportunity can be encountered more quickly. Between decision contexts, choices made in a fast speed-accuracy trade-off regime are usually followed by faster movements compared to those made in a regime encouraging slow and accurate choices (Thura et al., 2014; Thura, 2020). Altogether, these observations indicate that the level of urgency at which a decision is made directly influences movement vigor, suggesting that decision and movement durations are determined by a global decision urgency/movement vigor signal that invigorates behavior in order to control reward rate (Cisek and Thura, 2018; Carland et al., 2019). However, a missing test of the shared regulation hypothesis required to vary the motor context in which a decision is made and assess whether or not a motor context permitting execution of vigorous movements to express choices leads to faster decisions compared to the same difficult decisions made in a demanding motor context, imposing slow and accurate movements. Contrary to this prediction, we did not observe a robust and consistent effect of movement speed per se on decision duration and accuracy (by comparing short versus long target conditions, Figure 5B). Instead, data indicate that target size imposes a motor accuracy cost that is tackled by some subjects by shortening the deliberation period (Figure 5A) so that more time is available to prepare the following movement execution. This interpretation is supported by a post-experiment interview during which most of the participants declared having consciously expedited and thus “sacrificed” their decisions to better prepare action execution in small target trials.

One critical assumption in this experiment is that action towards the smaller targets requires less vigor compared to the action executed toward large targets. However, an alternative interpretation would state that because small targets impose more preparation time (reaction times are overall longer for small targets than for big targets in the DR task, figure 5C), a potential preparation-related urgency would have more time to increase in the small target blocks compared to the big target blocks, leading to movements initiated under higher urgency in the small blocks compare to the big blocks. The slower velocity and longer movement duration observed in the small blocks (Figure 4A for the tokens task) would then be explained by possible different systems for governing action preparation and execution (e.g. Haith et al., 2016). However, we do not believe that a putative preparation-related urgency signal could explain our results because no influence of this urgency is expected at the beginning of the trial and during the deliberation process. Instead, the preparation-related urgency level might differ between the blocks only after commitment, i.e. during movement preparation.

Thus, the present results more likely demonstrate that an unconditional and unidirectional relationship between action vigor and decision duration, as predicted by the shared regulation hypothesis, is absent. Instead, our results claim for a flexible mechanism in which decision and action durations are regulated by independent, yet interacting, decision urgency and movement vigor signals. Such flexibility is certainly advantageous given the inherent complexity of the many variables interrelationships at play during goal-directed behavior, where no single decision policy is guaranteed to maximize the reward rate across all contexts.

Flexibility between decision-making and action execution is well illustrated by the relationship between the effect of target size on decision duration in the tokens task and the effect of target size on reaction time in the delayed reach (DR) task. The significant correlation (Figure 6) indicates that subjects who are slower to initiate a movement in the small target trials of the DR task are also the subjects who adjust their decision policy the most in these difficult trials in the tokens task. The former result is consistent with data suggesting that effortful movements discount reward value, thus motivation, delaying the initiation of movements (Mazzoni et al., 2007; Summerside et al., 2018; Shadmehr et al., 2019). This relationship thus suggests that economic principles governing behavior utility in non-decision tasks extend to decision-making. It also indicates that when the task difficulty mainly relies on movement execution, as in the DR task, movement effort slows down reaction times whereas when task difficulty is shared between decision and action, as in the tokens task, movement effort influences the decision process in an opposite way. What could be the relevance of this intriguing behavior in terms of performance?

### Impact of a demanding movement on reward rate

The present data indicate that movement accuracy requirements, more than speed or duration, forced some subjects to hasten their decisions. It seems that they took advantage of the potentially long deliberation period permitted in the task (up to 3s) to sometimes shorten their judgment in order to focus on the following movement execution. Interestingly, such adjustment only occurred in blocks of trials in which decisions were encouraged to be conservative (“slow” decision blocks, Figure 7). Indeed, the large and very profitable, in terms of reward rate, shortening of decision durations observed in the “fast” decision blocks (Figure 8) probably constrained decision policy too much, preventing any other adjustments of behavior. It is also important to remember that in the tokens task, deciding more quickly does not provide additional time to execute the movement, the maximum movement duration being fixed at 800ms regardless of subjects’ reach onset timing. How then can one explain this suboptimal strategy? One possibility is that our limited cognitive and motor resources imposed a necessary trade-off between decision and action when task constraints were too demanding (Wickens, 2002). In this view, subjects had to choose between allocating resources on decision-making while taking the risk of producing inaccurate movements or rather sacrificing decision-making to presumably better prepare and execute their movements. Knowing that in ecological situations as in the present task, a movement usually follows the decision, it is possible that subjects gave priority to the action process considering that movement failure would prevent reward acquisition even if the decision was correct. Although it may be advantageous in terms of reward rate to decide very quickly while sacrificing a little bit of precision (see equation 3), as observed when humans and monkeys decide faster in the fast compared to the slow decision block of trials (Figure 8 and Thura et al., 2014; Thura, 2020), our results show however that the strategy consisting of sacrificing decision accuracy to execute accurate movements led to a drop of reward rate compared to a condition in which such adjustment was not necessary. This is probably because in small target trials, the probability of choosing the correct target decreased, even if the amount of time saved during the deliberation period compensated the longer movements made in this condition (Figure 4).

### Possible neurophysiological origin of the decision and action regulation mechanism

The interaction between the decision and action regulations provides a clue to the neural origins of the signals implicated in this mechanism. Interacting decision urgency and movement vigor signals would be expected to originate from a region that projects to a wide range of cortical areas to influence both decision-making and action execution. In this respect, the basal ganglia (BG) provide a natural candidate. The BG have long been functionally associated with the regulation of motivated behavior and reinforcement learning for maximizing reward (Graybiel, 2005; Frank, 2011), and multiple lines of neuropsychological, neurological and neurophysiological evidence suggest that effort expenditure and movement vigor are largely under the control of activity within a variety of BG structures, including the striatum, substantia nigra, ventral pallidum, and the globus pallidus (Mazzoni et al., 2007; Turner and Desmurget, 2010; Rueda-Orozco and Robbe, 2015; Dudman and Krakauer, 2016; Thura and Cisek, 2017; da Silva et al., 2018; Yttri and Dudman, 2018; Carland et al., 2019; Fobbs et al., 2020). All these studies along with results from the present report suggest a mechanism in which different populations of cells, located in the BG output nuclei, vary their activity to adjust both decision and motor durations under specific circumstances, in order to control the rate of reward. Future experiments designed to record the activity of individual BG cells during decision-making between actions in different decision and motor contexts should allow us to better understand the neural correlates of this regulation mechanism.

### Limitations

A limitation of the present study, as often in investigations of primate cognition and behavior, relates to the between-subject variability of the results. The average decision duration ranges from ∼700ms to about 1600ms depending on subjects (Figure 5), even though participants faced the same trials under identical conditions. This indicates individual “traits” of decision behavior. Similarly, a subgroup of four subjects was more vigorous than the others to execute their movements (Figure 4). While revealing probable unaddressed phenomena, these multiple levels of variability are still compatible with a flexible regulation mechanism of decision and action durations that would be idiosyncratic in nature. Another limitation concerns the absence of analysis of decision data in inaccurate or slow movement trials for methodology reasons. In the present report, we show that a difficult movement is often preceded by a fast and inaccurate decision, but this occurs when movements are properly executed. It is possible that subjects sometimes allocated their attention on the decision process, leading in that case to a “sacrifice” of motor control, resulting in failed movements. Further experiments or analyses are needed to reveal which of the two processes, the decision or the action, is typically prioritized by participants in this kind of demanding goal-directed behavior.

## ACKNOWLEDGMENTS

The authors wish to thank Sonia Alouche and Jean-Louis Borach for effective administrative assistance, Paul Cisek for his contribution in setting up the software environment, Frédéric Volland for his expertise during the technical preparation of this experiment, and Martine Meunier for helpful suggestions on the manuscript.

## REFERENCES

Balci F, Simen P, Niyogi R, Saxe A, Hughes JA, Holmes P, Cohen JD (2011) Acquisition of decision making criteria: reward rate ultimately beats accuracy. Atten Percept Psychophys 73:640–657.

Bogacz R, Hu PT, Holmes PJ, Cohen JD (2010) Do humans produce the speed–accuracy trade-off that maximizes reward rate? Quarterly Journal of Experimental Psychology 63:863–891.

Burk D, Ingram JN, Franklin DW, Shadlen MN, Wolpert DM (2014) Motor Effort Alters Changes of Mind in Sensorimotor Decision Making Kiebel S, ed. PLoS ONE 9:e92681.

Carland MA, Thura D, Cisek P (2019) The Urge to Decide and Act: Implications for Brain Function and Dysfunction. Neuroscientist:107385841984155.

Choi JES, Vaswani PA, Shadmehr R (2014) Vigor of Movements and the Cost of Time in Decision Making. Journal of Neuroscience 34:1212–1223.

Christopoulos V, Bonaiuto J, Andersen RA (2015) A biologically plausible computational theory for value integration and action selection in decisions with competing alternatives. PLoS Comput Biol 11:e1004104.

Christopoulos V, Schrater PR (2015) Dynamic Integration of Value Information into a Common Probability Currency as a Theory for Flexible Decision Making. PLoS Comput Biol 11:e1004402.

Churchland AK, Kiani R, Shadlen MN (2008) Decision-making with multiple alternatives. Nat Neurosci 11:693–702.

Cisek P, Puskas GA, El-Murr S (2009) Decisions in Changing Conditions: The Urgency-Gating Model. Journal of Neuroscience 29:11560–11571.

Cisek P, Thura D (2018) Neural circuits for action selection. In: Reach-to-grasp behavior: Brain, behavior, and modelling across the life span, Daniela Corbetta and Marco Santello., pp 91–118 Frontiers of developmental science. Taylor & Francis Group.

Cos I, Bélanger N, Cisek P (2011) The influence of predicted arm biomechanics on decision making. Journal of Neurophysiology 105:3022–3033.

Cos I, Duque J, Cisek P (2014) Rapid prediction of biomechanical costs during action decisions. Journal of Neurophysiology 112:1256–1266.

Cos I, Medleg F, Cisek P (2012) The modulatory influence of end-point controllability on decisions between actions. Journal of Neurophysiology 108:1764–1780.

da Silva JA, Tecuapetla F, Paixão V, Costa RM (2018) Dopamine neuron activity before action initiation gates and invigorates future movements. Nature 554:244–248.

Diamond JS, Wolpert DM, Flanagan JR (2017) Rapid target foraging with reach or gaze: The hand looks further ahead than the eye. PLoS Comput Biol 13:e1005504.

Ditterich J (2006) Evidence for time-variant decision making. European Journal of Neuroscience 24:3628–3641.

Dudman JT, Krakauer JW (2016) The basal ganglia: from motor commands to the control of vigor. Current Opinion in Neurobiology 37:158–166.

Fobbs WC, Bariselli S, Licholai JA, Miyazaki NL, Matikainen-Ankney BA, Creed MC, Kravitz AV (2020) Continuous Representations of Speed by Striatal Medium Spiny Neurons. J Neurosci 40:1679–1688.

Frank MJ (2011) Computational models of motivated action selection in corticostriatal circuits. Current Opinion in Neurobiology 21:381–386.

Gold JI, Shadlen MN (2007) The Neural Basis of Decision Making. Annu Rev Neurosci 30:535–574.

Graybiel AM (2005) The basal ganglia: learning new tricks and loving it. Current Opinion in Neurobiology 15:638–644.

Hagura N, Haggard P, Diedrichsen J (2017) Perceptual decisions are biased by the cost to act. eLife 6:e18422.

Haith AM, Pakpoor J, Krakauer JW (2016) Independence of Movement Preparation and Movement Initiation. J Neurosci 36:3007–3015.

Haith AM, Reppert TR, Shadmehr R (2012) Evidence for Hyperbolic Temporal Discounting of Reward in Control of Movements. Journal of Neuroscience 32:11727–11736.

Kawagoe R, Takikawa Y, Hikosaka O (1998) Expectation of reward modulates cognitive signals in the basal ganglia. Nat Neurosci 1:411–416.

Kira S, Yang T, Shadlen MN (2015) A Neural Implementation of Wald’s Sequential Probability Ratio Test. Neuron 85:861–873.

Malhotra G, Leslie DS, Ludwig CJH, Bogacz R (2017) Overcoming indecision by changing the decision boundary. Journal of Experimental Psychology: General 146:776–805.

Malhotra G, Leslie DS, Ludwig CJH, Bogacz R (2018) Time-varying decision boundaries: insights from optimality analysis. Psychon Bull Rev 25:971–996.

Marcos E, Cos I, Girard B, Verschure PFMJ (2015) Motor Cost Influences Perceptual Decisions Gribble PL, ed. PLoS ONE 10:e0144841.

Mazzoni P, Hristova A, Krakauer JW (2007) Why Don’t We Move Faster? Parkinson’s Disease, Movement Vigor, and Implicit Motivation. Journal of Neuroscience 27:7105–7116.

Michalski J, Green AM, Cisek P (2020) Reaching decisions during ongoing movements. Journal of Neurophysiology 123:1090–1102.

Morel P, Ulbrich P, Gail A (2017) What makes a reach movement effortful? Physical effort discounting supports common minimization principles in decision making and motor control Rushworth M, ed. PLoS Biol 15:e2001323.

Murphy PR, Boonstra E, Nieuwenhuis S (2016) Global gain modulation generates time-dependent urgency during perceptual choice in humans. Nat Commun 7:13526.

Myerson J, Green L (1995) Discounting of delayed rewards: models of individual choice. Journal of the Experimental Analysis of Behavior:263–276.

Neumann J von, Morgenstern O (1944) Theory of Games and Economic Behavior, Princeton University Press.

Reppert TR, Rigas I, Herzfeld DJ, Sedaghat-Nejad E, Komogortsev O, Shadmehr R (2018) Movement vigor as a traitlike attribute of individuality. Journal of Neurophysiology 120:741–757.

Revol P, Collette S, Boulot Z, Foncelle A, Niki C, Thura D, Imai A, Jacquin-Courtois S, Cabanac M, Osiurak F, Rossetti Y (2019) Thirst for Intention? Grasping a Glass Is a Thirst-Controlled Action. Front Psychol 10:1248.

Rueda-Orozco PE, Robbe D (2015) The striatum multiplexes contextual and kinematic information to constrain motor habits execution. Nat Neurosci 18:453–460.

Shadmehr R, Huang HJ, Ahmed AA (2016) A Representation of Effort in Decision-Making and Motor Control. Current Biology 26:1929–1934.

Shadmehr R, Orban de Xivry JJ, Xu-Wilson M, Shih T-Y (2010) Temporal Discounting of Reward and the Cost of Time in Motor Control. Journal of Neuroscience 30:10507–10516.

Shadmehr R, Reppert TR, Summerside EM, Yoon T, Ahmed AA (2019) Movement Vigor as a Reflection of Subjective Economic Utility. Trends in Neurosciences 42:323–336.

Standage D, You H, Wang D-H, Dorris MC (2011) Gain Modulation by an Urgency Signal Controls the Speed–Accuracy Trade-Off in a Network Model of a Cortical Decision Circuit. Front Comput Neurosci 5 Available at: http://journal.frontiersin.org/article/10.3389/fncom.2011.00007/abstract [Accessed September 9, 2019].

Steinemann NA, O’Connell RG, Kelly SP (2018) Decisions are expedited through multiple neural adjustments spanning the sensorimotor hierarchy. Nat Commun 9:3627.

Summerside EM, Shadmehr R, Ahmed AA (2018) Vigor of reaching movements: reward discounts the cost of effort. Journal of Neurophysiology 119:2347–2357.

Thura D (2020) Decision urgency invigorates movement in humans. Behavioural Brain Research 382:112477.

Thura D, Beauregard-Racine J, Fradet C-W, Cisek P (2012) Decision making by urgency gating: theory and experimental support. J Neurophysiol 108:2912–2930.

Thura D, Cisek P (2014) Deliberation and commitment in the premotor and primary motor cortex during dynamic decision making. Neuron 81:1401–1416.

Thura D, Cisek P (2016) Modulation of Premotor and Primary Motor Cortical Activity during Volitional Adjustments of Speed-Accuracy Trade-Offs. J Neurosci 36:938–956.

Thura D, Cisek P (2017) The Basal Ganglia Do Not Select Reach Targets but Control the Urgency of Commitment. Neuron 95:1160-1170.e5.

Thura D, Cos I, Trung J, Cisek P (2014) Context-dependent urgency influences speed-accuracy trade-offs in decision-making and movement execution. J Neurosci 34:16442–16454.

Todorov E, Jordan MI (2002) Optimal feedback control as a theory of motor coordination. Nat Neurosci 5:1226–1235.

Turner RS, Desmurget M (2010) Basal ganglia contributions to motor control: a vigorous tutor. Current Opinion in Neurobiology 20:704–716.

Wickens CD (2002) Multiple resources and performance prediction. Theoretical Issues in Ergonomics Science 3:159–177.

Wickler SJ, Hoyt DF, Cogger EA, Hirschbein MH (2000) Preferred speed and cost of transport. Journal of Experimental Biology:2195–2200.

Yoon T, Geary RB, Ahmed AA, Shadmehr R (2018) Control of movement vigor and decision making during foraging. Proc Natl Acad Sci USA 115:E10476–E10485.

Yttri EA, Dudman JT (2018) A Proposed Circuit Computation in Basal Ganglia: History-Dependent Gain: Proposed Circuit Computation in Basal Ganglia. Mov Disord 33:704–716.

